# A global serosurvey of RNA virus-reactive antibodies in bats and rodents reveals the enzootic presence of flavi- and paramyxoviruses in African Pteropodidae

**DOI:** 10.1101/2025.10.28.684997

**Authors:** Jackson Emanuel, Jan Papies, Leonie Meiners, Talitha Veith, Victor M. Corman, Tabea Binger, Erik Lattwein, Beate M. Kümmerer, Susanne Zweerink, Kai Fechner, Florian Gloza-Rausch, Veronika Cottontail, Gaël D. Maganga, Jonas Schmidt-Chanasit, Peter Vallo, Samuel K. Oppong, Marco Tschapka, Chantal B. E. M. Reusken, Rainer G. Ulrich, Eric M. Leroy, Jan Felix Drexler, Terry C. Jones, Christian Drosten, Marcel A. Müller

## Abstract

Bats and rodents have been identified as reservoir hosts of diverse RNA viruses that pose potential risks to humans. However, overall virus prevalence in wildlife and underlying factors determining the zoonotic potential of reservoir-borne viruses remain poorly characterized. Virus detection often relies on the identification of viral nucleotide sequences and may therefore be limited by variable virus concentration, tissue tropism, sample quality, and duration of infection. In this study, we applied a multiplex immunofluorescence assay to examine bat and rodent seroprevalence against 14 different human pathogenic RNA viruses spanning seven virus families (*Coronaviridae, Flaviviridae, Hantaviridae, Paramyxoviridae, Phenuiviridae, Pneumoviridae, Togaviridae*). We retrospectively analyzed 1,135 bat (16 species) and 454 rodent (15 species) blood or transudate specimens collected at 63 sites in seven countries. Tropical bats exhibited notably high rates of seropositivity against certain virus families (*Flaviviridae:* 42.8%; *Paramyxoviridae:* 60.4%; *Togaviridae*: 7.5%). Multivariable logistic regression models were created to evaluate the association of tropical bat characteristics with seropositivity. Preliminary findings indicate that large colony size, found among species such as *Eidolon helvum* and *Rousettus aegyptiacus*, is a risk factor for orthoflavi- and orthorubulavirus infection, consistent with the communal spread of bat-adapted viruses.

**Author Summary:** Many animal populations carry viruses that have the potential to spill over to humans and cause severe disease. Despite surveillance efforts, it is logistically challenging to detect virus infections in wildlife, as viruses may have a short duration of infection or unknown routes of transmission. In this study, we examined blood and transudate samples from rodents and bats around the world for antibodies against 14 different viruses known to cause disease in humans. We analyzed the samples via a mosaic chip-based immunofluorescence test, which has the advantage of detecting antibodies against whole-virus antigen. This method allows for broad surveillance, as we would also expect to uncover evidence of past infection with bat- and rodent-adapted viruses that are related, but not identical, to known human pathogens. We detected antibodies against all seven virus families we examined, with especially high detection rates of antibodies in tropical bats. We used the results to construct multivariable logistic regression models of tropical bat seroprevalence to examine potential associations between tropical bat characteristics and previous infection by certain virus families. Bats with large colony sizes, such as *Eidolon helvum* and *Rousettus aegyptiacus*, were apparently at increased risk of infection with orthoflavi- and orthorubulaviruses compared to other tropical bat species.

## Introduction

Zoonotic viruses emerging from wildlife reservoirs continue to pose a major threat to human health [1–3]. It has been estimated that there are approximately 10,000 potentially zoonotic viruses with mammalian hosts, emphasizing the need for broad surveillance [4]. While the sources of zoonotic viruses are diverse, bats (order Chiroptera) and rodents (order Rodentia) host a wide range of enzootic viruses that are co-ancestral with prominent human pathogens. Examples include coronaviruses [5, 6], filoviruses [7], lyssaviruses [8], and paramyxoviruses [9] in bats, as well as arenaviruses [10] and hantaviruses [11] in rodents. It has recently been proposed that the proportion of zoonotic viruses capable of infecting humans is consistent among mammalian taxonomic orders [12]. This suggests that the relatively large number of zoonotic viruses originating from bats and rodents may be a function of high species richness in these orders [12, 13]. In any case, virus surveillance efforts often focus on a small number of virus families or host species, leaving open the possibility that remote areas or undersampled species with high virus prevalence remain undetected. Better identification of virus hotspots in nature may lead to targeted measures to reduce the risk of zoonotic spillover and contribute to “primary pandemic prevention” [14].

In addition to identifying host species with high seroprevalence, determining the underlying factors associated with increased infection risk is of considerable scientific value. By analyzing behavioral and life history traits among diverse taxa, statistical models may be employed to identify associations with seropositivity. This has the advantage of potentially providing evidence of previously unknown routes of transmission. For example, some arthropod-borne viruses (arboviruses), including flavi- and togaviruses, appear to maintain sylvatic transmission cycles in non-human primates [15], but potential involvement of other mammalian host species and insect-independent transmission (e.g., droplet or fecal-oral) have not been excluded. Indeed, some flaviviruses with no known vector, such as Entebbe bat virus, have been exclusively isolated from bat tissue, including salivary glands, raising the possibility of arthropod-independent transmission cycles [16, 17]. Systematic investigation of infection rates in bats and rodents might help discern between enzootic and epizootic hosts and reveal unknown maintenance cycles amenable to control measures.

There are several ways to assess pathogen diversity in putative hosts, such as PCR-based techniques, high-throughput sequencing, and virus isolation using cell culture. All methods that involve direct virus detection require samples from periods of active infection. This may be challenging due to spatiotemporal variability in virus prevalence, unknown virus tissue tropism, and field conditions which result in nucleic acid degradation or loss of infectivity. By contrast, examining the seroprevalence (i.e. virus-specific antibodies) of host populations provides evidence that individuals have previously been exposed to a pathogen, regardless of acute infection status. Moreover, by using whole-virus antigens, one can screen for groups of related viruses that exhibit cross-reactivity due to shared antigenic properties. Even virus families which have been grouped into distinct serotypes may exhibit such cross-reactivity [18]. Serological approaches have previously led to the discovery of novel hepaciviruses in horses and rodents [19, 20], confirmed the presence of severe acute respiratory syndrome coronavirus (SARS-CoV)-like antigen in African bats [21], and revealed exposure with filovirus antigens in bats and human primates in non-endemic regions of Africa [22, 23].

In this study, we assessed the presence of antibodies against RNA viruses in a large sample of sera and peritoneal transudates from bats and rodents collected in tropical and temperate climate zones. Serology was based on immunofluorescence tests using whole-virus antigens from 14 different human pathogenic RNA virus species, representing seven different viral families. Our findings indicated a high seroprevalence among representatives of tropical bat genera, which prompted us to construct multivariable logistic regression models to evaluate tropical bat host characteristics as potential risk factors for infection. Although results from these analyses were challenged by the need to disambiguate host characteristics from other potentially confounding factors, we report our findings and provide a framework for future modeling efforts.

## Methods

### Patient serum samples characterized by indirect immunofluorescence test (IIFT)

As positive controls and to evaluate cross-reactivity, human serum samples (n=421) were obtained from the EUROIMMUN biobank and tested by EUROIMMUN. Specifically, reference control sera were defined as either having derived from patients after acute infection or vaccination with the virus of interest or having dominant titers (i.e., at least tenfold higher against the virus of interest compared to related viruses). Control sera exhibiting reactivity against a variety of human pathogens were used, including West Nile virus (WNV, *Orthoflavivirus nilense*, n=23), yellow fever virus (YFV, *Orthoflavivirus flavi*, n=33), Dengue virus serotype 2 (DENV2, *Orthoflavivirus denguei*, n=22), tick-borne encephalitis virus (TBEV, *Orthoflavivirus encephalitidis*, n=32), Japanese encephalitis virus (JEV, *Orthoflavivirus japonicum*, n=24), hepatitis C virus (HCV, *Hepacivirus hominis*, n=20), *Orthohantavirus* spp. (n=21), Crimean-Congo hemorrhagic fever virus (CCHFV, *Orthonairovirus haemorrhagiae*, n=18), measles virus (MeV, *Morbillivirus hominis*, n=33), mumps virus (MuV, *Orthorubulavirus parotidis*, n=25), respiratory syncytial virus (RSV, *Orthopneumovirus hominis*, n=32), human parainfluenzavirus 1 (PIV1, *Respirovirus laryngotracheitidis*, n=29), human parainfluenzavirus 2 (PIV2, *Orthorubulavirus laryngotracheitidis*, n=32), human parainfluenzavirus 3 (PIV3, *Respirovirus pneumoniae*, n=33), human parainfluenzavirus 4 (PIV4, *Orthorubulavirus hominis*, n=33), Chikungunya virus (CHIKV, *Alphavirus chikungunya*, n=9), and Sindbis virus (SINV, *Alphavirus sindbis*, n=2). Reference sera were tested via indirect immunofluorescence tests (IIFT) against relevant subsets of the full mosaic panel. All serum samples were prepared according to the manufacturer’s instructions for the respective IIFT product (EUROIMMUN AG, Lübeck, Germany).

### Animal sampling

Bat (n=1,135) and rodent (n=454) serum samples were collected from 2005 through 2011 in seven different countries, i.e. Germany, the Netherlands, Gabon, Ghana, the Republic of Congo, Thailand, and Panama (**Fig. 1**). Bat samples comprised 16 species from 14 genera and rodent samples comprised 15 species from 5 genera (**Fig. 1**; **Table 1**). As previously described, all animals were caught and identified morphologically by trained field biologists [9, 20, 24]. All sampling was approved by the responsible animal ethics committees and all efforts were made to minimize the suffering of animals, including the application of sodium pentobarbital/ketamine anesthesia when surgical procedures were performed.

**Fig. 1.**
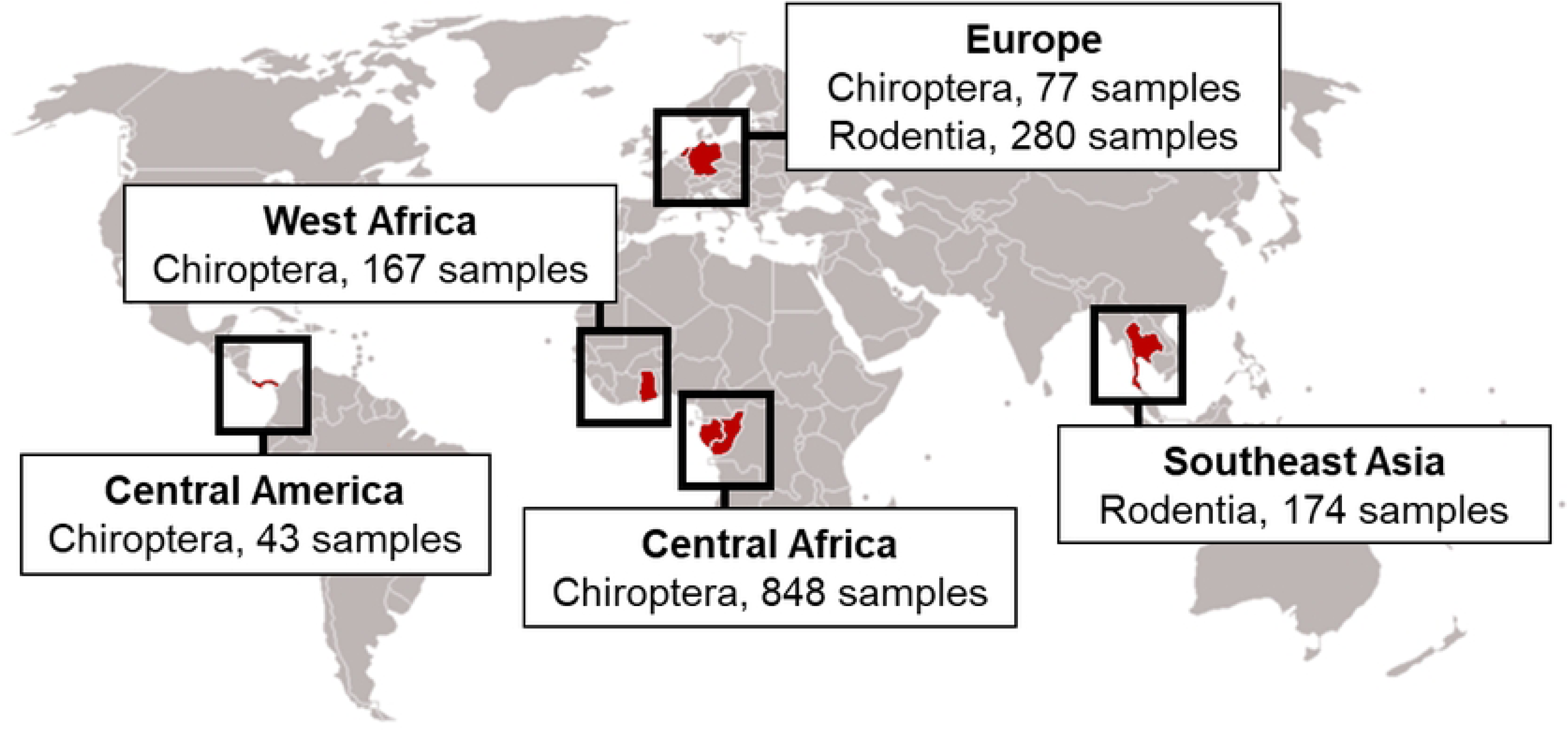
Overview of sampling sites. Samples were obtained from 63 sites, encompassing five geographic regions: Central America (Panama), Europe (Netherlands, Germany), West Africa (Ghana), Central Africa (Gabon, Republic of Congo), and Southeast Asia (Thailand). Serum and transudate samples were collected from 2005 through 2011. Bat samples comprised 16 species from 14 genera; rodent samples comprised 15 species from 5 genera. This map depicts the total number of serum and transudate samples obtained from each region.

**Table 1.**
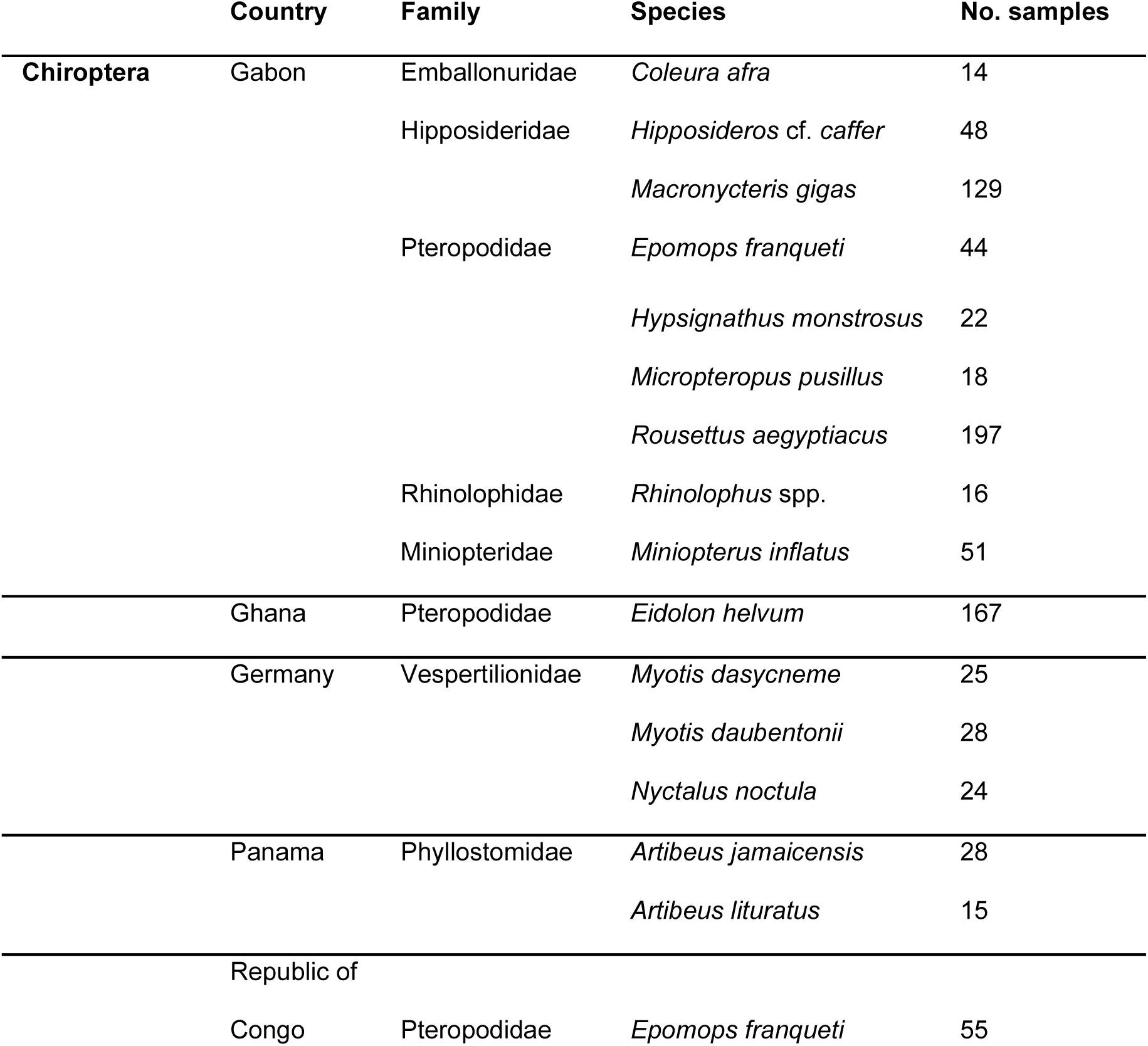
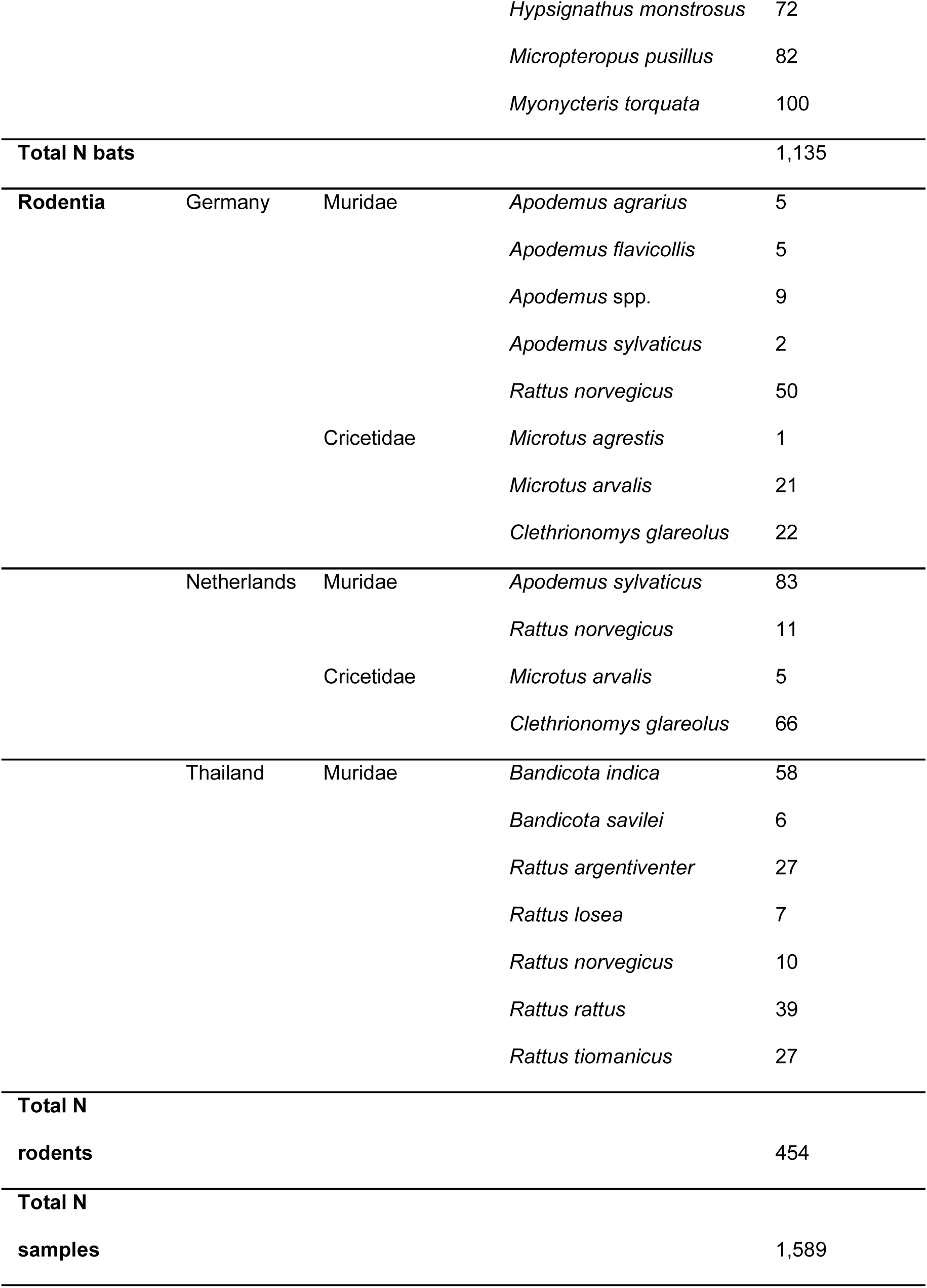
Overview of analyzed bat and rodent samples.

### Immunoglobulin species-reactivity control by enzyme-linked immunosorbent assay (ELISA)

As we tested bats and rodents from different species, recognition of individual immunoglobulins (Ig) by the applied secondary IgG, i.e. goat-anti bat or goat-anti mouse Ig, was tested. Serum samples (two from each species) were applied in duplicate in an enzyme-linked immunosorbent assay (ELISA) as described elsewhere [25, 26]. Briefly, serum specimens were diluted 1:200 in PBS-Tween/1% milk powder and coated on Maxisorb ELISA plates at 4°C overnight. Plates were blocked with PBS-T/5% milk powder and detection was achieved by first incubating with goat anti-bat or goat anti-mouse IgG (Bethyl Laboratories, Montgomery, USA) diluted 1:1,000 in PBS-T/1% milk powder followed by a horseradish peroxidase-labeled donkey anti-goat IgG (Dianova, Hamburg, Germany) diluted 1:8,000 in PBS-T/1% milk powder. Substrate TMB (Mikrogen, Neuried, Germany) was added and the ELISA plates were read at 450/630 nm. Background signals (incubation with second and third antibody only) were subtracted and all optical density values were multiplied by 1,000.

### Screening by mosaic chip-based indirect immunofluorescence test (mIIFT)

For screening of serum specimens, we used a custom-designed mosaic chip-based indirect immunofluorescence test (mIIFT) with minor modifications as described previously [27, 28]. The assay allowed to simultaneously screen for antibody reactivity against 14 viruses from seven different families (*Coronaviridae, Flaviviridae, Hantaviridae, Paramyxoviridae, Phenuiviridae, Pneumoviridae, Togaviridae*). To evaluate intra-assay cross-reactivity with related viruses, a variety of prototypic viruses were selected: human coronavirus 229E (HCoV-229E), SARS-CoV, West Nile Fever virus (WNV), yellow fever virus (YFV), Dengue virus serotype 2 (DENV2), Measles morbillivirus (MeV), respiratory syncytial virus (RSV), human parainfluenzavirus 1 (PIV1), human parainfluenzavirus 2 (PIV2), mumps virus (MuV), Dobrava-Belgrade orthohantavirus (DOBV), Rift Valley fever virus (RVFV), Chikungunya virus (CHIKV), and Venezuelan equine encephalitis virus (VEEV). All serum samples were diluted at 1:40 in sample buffer (EUROIMMUN AG, Lübeck, Germany) and incubated for 2 hours at room temperature. After washing with phosphate-buffered saline (PBS) including 0.1% Tween, serum-derived antibody binding was detected with a goat anti-bat (Bethyl, Montgomery, AL, USA) or goat anti-mouse IgG (Dianova) diluted 1:1,000 and a Dylight488 labeled donkey anti-goat IgG (1:100). Analysis was performed with a Motic Immunofluorescence microscope (Zeiss, Jena, Germany). For comparison, all pictures were taken with identical microscope settings.

### Virus neutralization assays

Neutralizing activities of bat sera against YFV were tested by plaque reduction assay on BHK-21/J cells (kindly provided by Charles Rice, The Rockefeller University, NY, USA). 50 plaque-forming units of YFV were mixed with different dilutions of heat-inactivated bat serum (ranging from 1:10 to 1:320) and incubated for 1 h at 37°C. Virus-serum mixtures were then added to BHK-21/J cells seeded in 24-well plates. Following incubation for 1 h at 37°C, the medium was replaced with an agarose overlay (MEM with 0.6% agarose and 2% FCS). After incubation at 37°C for 3 days, cells were fixed with 7% formaldehyde and stained with 1% crystal violet in 5% ethanol. Infectivity was calculated according to the equation [% infectivity = 100 x (plaques sero-neutralization sample / plaques virus infection sample)]. In the case of MuV, a TCID50-based neutralization test was performed in a 96-well plate using Vero E6 cells (ATCC CRL-1586). Sera and virus were diluted and incubated as described above and incubated for 1 h at 37°C. The inoculum was replaced with fresh medium and plates were incubated at 37°C for 10 days before fixation and staining.

### Statistical methods

The data obtained from tropical bats were selected for more extensive analysis, as both sample size (n=1,058) and seroprevalence (874/1058 samples positive for at least one virus) were sufficiently high to consider modeling risk factors associated with host seropositivity. Temperate bat sample size (n=77) and seroprevalence (29/77 samples were positive for at least one virus) were considerably more limited. As modes and dynamics of virus transmission may considerably differ between climate zones, and due to the small number of temperate bat samples with seroreactivity that could potentially inform such differences, we did not include data from temperate bats in the statistical analysis. The limited number of rodent genera and relatively low seroprevalence in this study was not conducive to generating analogous models for rodent seropositivity.

Using the tropical bat data, logistic regression analyses were performed separately for each virus species, with detected seropositivity as the binary outcome and the same set of input variables in each virus-specific regression model. Binary input variables were adult (as opposed to juvenile), cave-dwelling (as opposed to tree-dwelling), insectivorous diet (as opposed to frugivorous diet), migratory lifestyle (as opposed to sedentary lifestyle), sex (male as opposed to female), and membership in the taxonomic suborder Yangochiroptera (as opposed to Yinpterochiroptera). Expected lifespan and colony size (log-transformed) were added as continuous input variables. Values of all mentioned variables except for adult and sex were inferred from the sample’s assigned bat species. Two models were implemented, following the implications of a directed acyclic graph (DAG, **S1 Fig.**): model A included all aforementioned variables except lifespan as its inclusion led to problems with model convergence; model B contained additional categorical input variables: bat family, region of sampling (Ghana, Gabon/Republic of Congo, Panama), and a newly-created variable composed of the year and region of sampling, henceforth referred to as year_region. Unlike model A, which did not include potential confounders, model B contained collinear input variables increasing the uncertainty around parameter estimates and the sensitivity to initial conditions during sampling. For example, many bat families were only sampled in one region, and within bat families, often data from only one species exhibiting one specific combination of host characteristics is available. This should be borne in mind when interpreting the statistical results.

The prior distribution for the parameter corresponding to year_region was a Normal distribution with a mean of 0 and a standard deviation of 0.25 (N(0, 0.25)). For all remaining binary and categorical variables, encoded as indicator and index variables, respectively, prior distributions were set to N(0, 0.5). Region was modeled as a fixed effect while the remaining categorical variables were modeled as random effects (non-centered parameterization). Prior distributions for lifespan and log colony size parameters were set to N(0, 0.05) and N(0, 0.2), respectively.

We obtained estimates of and uncertainty around bat species lifespans and colony sizes from literature sources (**S1 Table**). To incorporate this information, “true” species lifespans and colony sizes were sampled from Log-Normal and truncated Normal distributions, respectively, using the species-specific means and extreme (minimum and maximum) values inferred from the literature. If no estimate of the mean lifespan was available for a species, the value from a species of the same genus or family was used. If no information was available, a value of 12 years was assigned. For *Rhinolophus* spp. obtained in Gabon, biological data from *Rhinolophus landeri* was used. For *Hipposideros* cf. *caffer* and *Macronycteris gigas,* we used the same lifespan expectancy as for *Rhinolophus* spp., as the corresponding families are closely related. Following manual exploration of implied Log-Normal distributions using mean life expectancy values with different values of sigma, the following sigma values were set: 0.2 if a lifespan estimate was available for the respective species, 0.3 if a lifespan estimate was available for a species from the same genus, 0.4 if a lifespan estimate was available for a species from the same family, and 0.6 if no lifespan information was available. Colony sizes were allowed to vary between different combinations of sampling year, sampling site, and sampled species. Note that this implementation assumes separate colony sizes for each species, even when samples were taken at the same site in the same year. The sampled value, along with the values of the remaining variables, were then used as input variables for the computation of the Bernoulli likelihood.

Model building and Markov Chain Monte Carlo (MCMC) sampling was performed with PyMC (version 5.10.0) using the BlackJAX NUTS sampler. Posterior analyses and plotting were done using ArviZ, version 0.16.1. Sampling was done for 4,000 iterations with 1,000 tuning steps. The following acceptance rates were used: 90% in all regression analyses using model A, 99% in the analysis of RVFV seropositivity using model B, 98% in the analyses of CHIKV, VEEV and SARS-CoV seropositivity using model B and 95% in the analyses of seropositivity of the remaining viruses using model B. Model convergence was verified using the Gelman Rubin statistic (Rhat) and visual inspection of posterior traces. Model fit was assessed with posterior predictive checks. Visualizations of seroprevalence based on species, sampling site, and sampling year, as well as source code and input data for the logistic regression models are all available at the following site: https://github.com/VirologyCharite/batSerology.

## Results

### Evaluating specificity of the IIFT assay against whole virus antigens using human control sera

Before screening animal sera, serological cross-reactivity was evaluated against a panel of human serum samples (n=421). Specifically, control samples that were known to be seropositive against a variety of human pathogens (including WNV, YFV, DENV2, TBEV, JEV, HCV, *Orthohantavirus*, CCHFV, MeV, MuV, RSV, PIV1, PIV2, PIV3, PIV4, CHIKV, and SINV) were tested by chip-based indirect immunofluorescence tests (IIFT) for cross-reactivity against relevant subsets of the mosaic panel. Cross-reactivity between SARS-CoV and HCoV-229E has been evaluated elsewhere [29] and was therefore excluded before analysis. Remaining viruses were clustered into four groups: *Flaviviridae* (WNV, YFV, DENV2), *Bunyaviricetes* (DOBV, RVFV), *Mononegavirales* (MeV, MuV, RSV, PIV1, PIV2), and *Togaviridae* (VEEV, CHIKV) whole-virus antigens. Within each of these four groups cross-reactivity was evaluated (**S2 Table**). For the flaviviruses, the strongest cross-reactivity was between WNV and DENV2, where 100% of WNV-positive control sera exhibited cross-reactivity against DENV2 antigen and 93.8% of DENV2-positive control sera exhibited cross-reactivity against WNV antigen. WNV- and DENV2-positive sera also exhibited strong reactivity against YFV antigen, 73.9% and 87.0% respectively. Only hepatitis C virus (HCV)-positive sera did not exhibit cross-reactivity with the other flavivirus antigens tested, suggesting that the orthoflaviviruses are serologically distinct from the hepaciviruses (**S2 Table**). In contrast to the flaviviruses, the *Bunyaviricetes* members, exhibited minimal cross-reactivity, with 5.9% of CCHFV-positive sera testing positive against RVFV, and no cross-reactivity detected between CCHFV and DOBV or *Orthohantavirus* and RVFV (**S2 Table**). Among *Mononegavirales* members, reference sera positive for one of the pneumo-or paramyxoviruses (MeV, MuV, RSV, PIV1, PIV2, PIV3, and PIV4) were often positive for all pneumo-or paramyxoviruses in the mIIFT (MeV, MuV, RSV, PIV1, PIV2). These rates ranged from 70.0% of PIV2-positive reference sera exhibiting reactivity against MuV in the IIFT to 97.0% of PIV3- and PIV4-positive reference sera exhibiting reactivity against PIV2 in the IIFT (**S2 Table**). Finally, the togaviruses also showed evidence of apparent cross-reactivity, with 6/9 CHIKV-positive reference sera exhibiting reactivity against VEEV and 1/2 SINV-positive reference sera exhibiting reactivity against CHIKV (**S2 Table**). It should be noted that these results do not explicitly provide evidence of cross-reactivity, particularly regarding the paramyxoviruses, as it is expected that most human reference sera are IgG-positive against both MeV and MuV due to high rates of vaccination. Nevertheless, these results are provided to better contextualize IgG IIFT performance on clinical samples with a known infection history.

### Evaluating the reactivity of goat anti-bat IgG and goat anti-mouse IgG antibodies against serum samples derived from various species

As we applied sera from multiple bat and rodent species, the reactivity of goat anti-bat and goat anti-mouse IgG conjugates was tested for each bat and rodent species used in this study. Animal sera were coated on ELISA plates and incubated with conjugate at a fixed dilution. Out of the 16 bat species, twelve exhibited comparable levels of reactivity with >0.1 OD_450-630_ (**S2 Fig.**). Four species (*Artibeus jamaicensis*, *Artibeus lituratus*, *Myotis dasycneme,* and *Nyctalus noctula*) showed less intense reactivity in the ELISA (**S2 Fig. a**). Panels of sera from these four species were used in initial experiments of virus antibody detection in mIIFT, indicating that fluorescence signal intensity in virus-infected cells was sufficiently high for the inclusion of these bat species in the study. For example, the low ELISA-reactive serum from *Artibeus jamaicensis* (P1018) showed a comparable or more pronounced and specific fluorescence signal for mumps and measles virus antigens when compared to a high ELISA-reactive serum from *Rousettus aegyptiacus* (GB687) (**S3 Fig.**). For rodents, sera from all tested species were recognized by the goat anti-mouse IgG with comparable efficiency (**S2 Fig. b**). Sampling efforts resulted in serum collection from the species summarized in **Fig. 1** and **Table 1**.

### Results from mIIFT assay screening of bat and rodent samples and virus neutralization assays

To screen for antibodies arising from virus infection, all samples were tested by mIIFT at a serum dilution of 1:40. A representative example of a reactive serum from *Rousettus aegyptiacus* in comparison to positive human control sera is presented in **S4 Fig.** The bat serum sample strongly recognized all three orthoflavivirus antigens (WNV, YFV, DENV2), both orthorubulaviruses (MuV and PIV2), and the RVFV antigen. Reactivity was rated from very high reactivity (+++) to negative (-) according to fluorescence signal intensity with fixed microscope settings. As sera originating from different species resulted in somewhat variable degrees of background fluorescence, reactivity was evaluated with respect to negative baselines for each species. An example of three different fluorescence intensity categories is given in **S5 Fig.** Sera with high background signal or inconclusive results were rated negative.

To further assess the specificity of the observed mIIFT reactivities, we randomly chose five MuV and five YFV mIIFT-positive and mIIFT-negative bat sera with variable cross-reactivity patterns for related viruses to perform highly specific virus neutralization tests (**Table 2**). Whereas 0/10 MuV mIIFT-positive sera were able to neutralize MuV, 1/10 YFV mIIFT-positive sera had a neutralizing antibody titer of 1:20 against YFV. Thus, mIIFT-positive samples did not necessarily exhibit neutralizing activity, which may be explained by cross-reactive antibodies resulting from previous infections with bat-specific viruses rather than the human-specific viruses used in the neutralization assay.

**Table 2.**
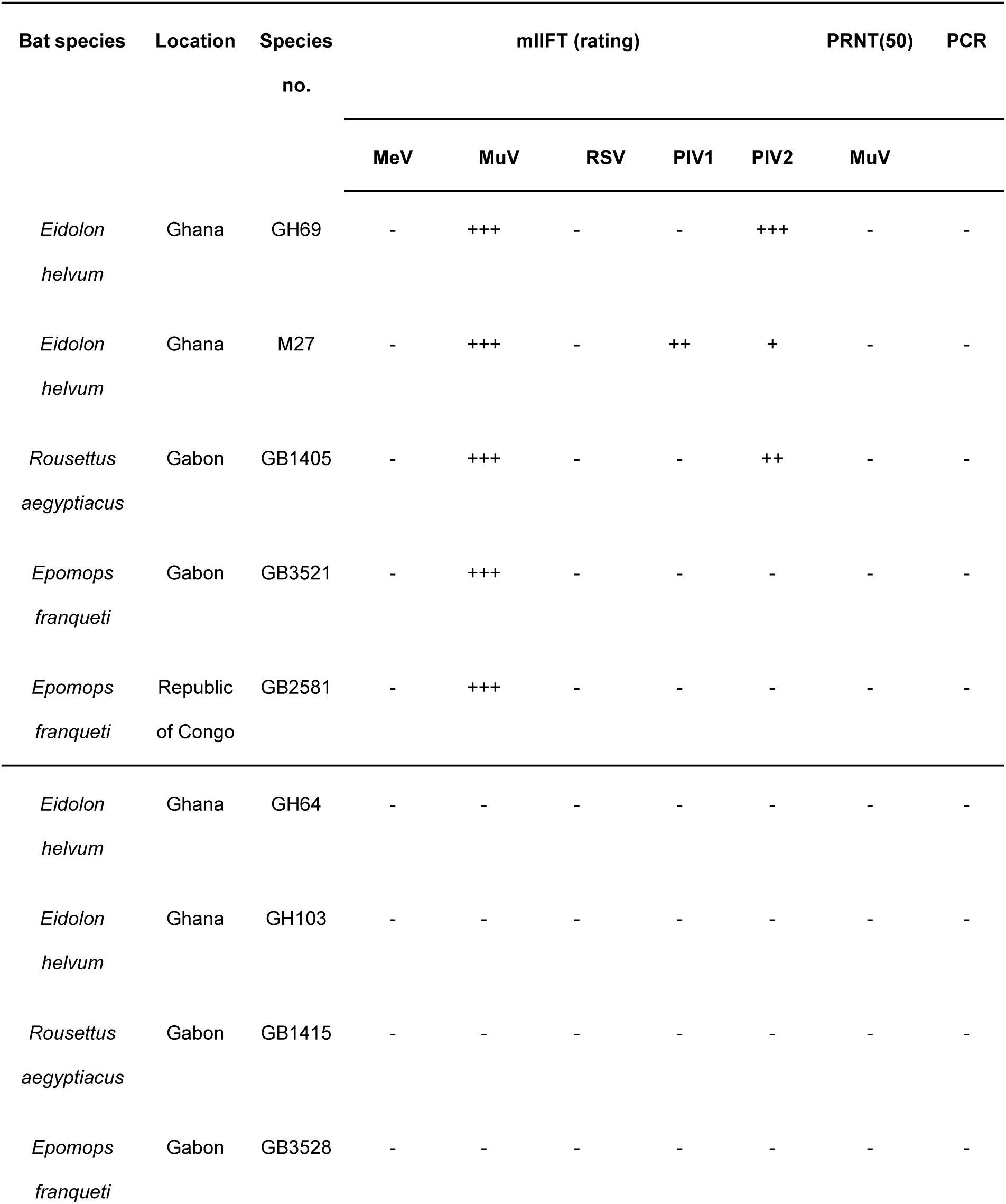
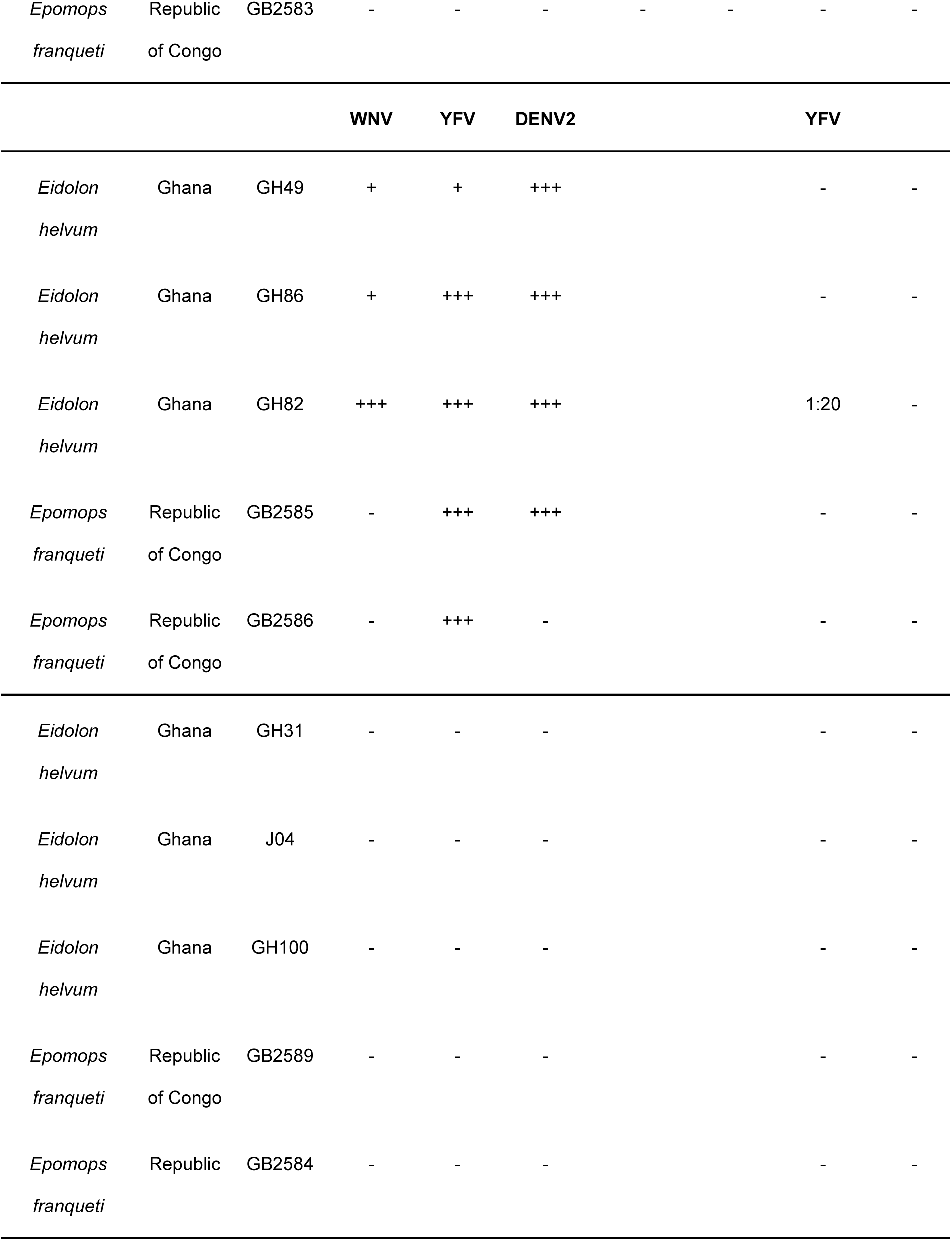
Mumps and Yellow fever virus plaque reduction neutralization assay for mIIFT-positive and mIIFT-negative bat sera.

As multiple samples were seropositive for more than one virus family, a complete overview of the seropositivity data is provided for all bat (**Fig. 2a**) and rodent (**Fig. 2b**) genera. Among bats, 905/1135 (79.7%) samples were seropositive against at least one virus family and 515/1135 (45.4%) were seropositive against two or more virus families. 34/1135 (3.0%) bat samples were seropositive against four or more virus families, indicating that at least a small fraction of individuals are infected with numerous, diverse viruses over the course of their lifespan. 821/950 (86.4%) Yinpterochiroptera samples were seropositive, whereas only 84/185 (54.6%) Yangochiroptera samples were seropositive. This apparent difference is partially a reflection of the fact that the temperate bat genera we examined, *Myotis* and *Nyctalus*, both Yangochiroptera, exhibited a lower rate of seropositivity 29/77 (37.7%). In particular, African bats, which constituted the majority of our samples (n=1015), exhibited notably high rates of seropositivity, with 860/1015 (84.7%) samples testing positive against at least one virus family. Even among the tropical bats, two species of interest, *Eidolon helvum* and *Rousettus aegyptiacus*, were notable for having an especially high rate of seropositivity against multiple virus families, with 140/197 (71.1%) and 136/167 (81.4%) seropositive against two or more virus families, respectively. Among rodents, 182/454 (40.1%) of samples were seropositive against at least one virus family and 45/454 (9.9%) were seropositive against two or more virus families. 280/454 (61.7%) rodent samples we analyzed were from a temperate climate zone, which may have contributed to relatively low rates of seropositivity in this dataset, as the three genera sampled exclusively in a temperate zone, *Apodemus* (29/104; 27.9%), *Clethrionomys* (22/88; 25.0%), and *Microtus* (8/27; 29.6%), had lower rates of seropositivity than the genus exclusively sampled in a tropical zone, *Bandicota* (38/64; 59.4%). Samples from the genus *Rattus* were collected in both temperate and tropical zones, with a higher rate of seropositivity observed in the tropical region (61/110; 55.5%) compared to the temperate region (24/61; 39.3%).

**Fig. 2.**
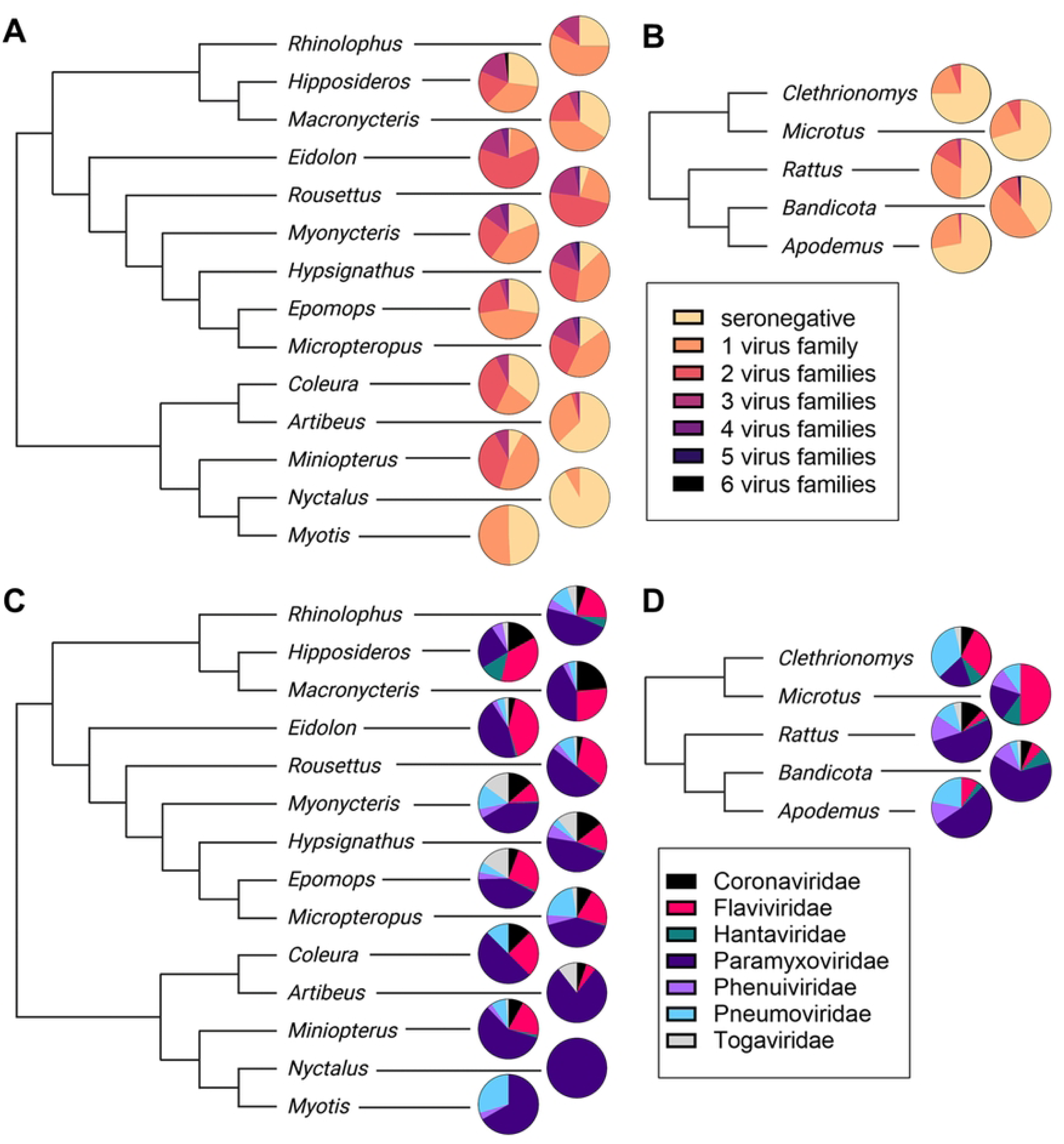
Graphical summary of seropositivity among bat and rodent genera. Samples included 19 mammalian genera (14 bat, 5 rodent). Dendrogram representations of their phylogenetic relationships are depicted. Because many samples were mIIFT-positive against more than one virus family, pie charts showing the fraction of samples positive for a given number of viral families were generated **(a)** for bats, and **(b)** for rodents. To visualize the seroprevalence of viral families among genera, pie charts showing the fraction of seropositive detection events for each viral family out of the total number of positive detection events **(c)** in bats, and **(d)** in rodents.

A comparison of virus family prevalence among seropositive samples is also provided for all bats (**Fig. 2c**) and all rodents (**Fig. 2d**). Among bats, two virus families were especially notable for having high rates of overall seropositivity. 736/1135 (64.9%) bat samples were seropositive against one or more paramyxoviruses and 452/1135 (39.8%) bat samples were seropositive against one or more flaviviruses. These rates were even higher among particular species, with 152/167 (91.0%) *Eidolon helvum* samples and 184/197 (93.4%) *Rousettus aegyptiacus* samples seropositive against one or more paramyxoviruses, and 131/197 (66.5%) *R. aegyptiacus* samples and 141/167 (84.4%) *E. helvum* samples seropositive against one or more flaviviruses. Seropositivity against other virus families, including the *Coronaviridae, Hantaviridae, Phenuiviridae, Pneumoviridae,* and *Togaviridae* was broadly present at low rates across many bat genera, consistent with sporadic, epizootic infection. Temperate bat genera, *Myotis* and *Nyctalus*, exhibited markedly less virus diversity than tropical bats, predominantly showing reactivity against paramyxoviruses and pneumoviruses. Among rodents, paramyxoviruses constituted a majority of positive detection events in *Apodemus* (17/32; 53.1%), *Bandicota* (31/49; 63.3%), and *Rattus* (61/117; 52.1%) whereas pneumoviruses and flaviviruses constituted a plurality of positive detection events in *Clethrionomys* (9/27; 33.3%) and *Microtus* (5/10; 50.0%), respectively. Seroprevalence against togaviruses was largely restricted to tropical animals, especially *Hypsignathus*, *Myonycteris*, and *Epomops* bats, as only a single temperate climate-dwelling rodent sample (*Rattus norvegicus* in Stuttgart, Germany) tested positive against a togavirus. Overall, the results in **Fig. 2** indicate widespread seroconversion of wild bats and rodents to diverse viruses closely related to known human pathogens. Due to sampling constraints and the presence of confounding factors, general comparisons about differences in seroprevalence between bats and rodents or between tropical and temperate climate zones are not explicitly supported by our dataset. The complete dataset is reported in **S3 Table**.

### Statistical modeling of tropical bat host characteristics associated with seropositivity

Two multivariable logistic regression models were used to identify tropical bat host characteristics associated with seropositivity. Model A, which was designed to limit multicollinearity, identified posterior density intervals for the following variables: adult/juvenile, cave-dwelling/tree-dwelling, insectivorous/frugivorous, migratory/non-migratory, male/female, Yangochiroptera/Yinpterochiroptera, and estimated mean colony size (**S6 Fig.**). Model B, which was designed to distinguish between potentially confounding factors, also identified posterior density intervals for the aforementioned variables as well as bat family, region of sampling (Ghana, Gabon/Republic of Congo, Panama), a variable composed of both the year and region of sampling, year_region, and bat species lifespan (**S7 Fig.**). Posterior density intervals from both models are reported for all viruses, except DOBV, for which no model could be generated due to a low rate of seropositivity, and PIV1 and RSV, for which only model A parameters are available due to poor model convergence when using model B.

Broadly speaking, Model A resulted in similar highest posterior density intervals (HPDIs) of expected change in log-odds for viruses of the same family, potentially indicative of cross-reactivity or of conserved virus family host preferences. Perhaps most strikingly, orthoflavivirus (DENV2, WNV, YFV) seropositivity showed a positive association with host frugivory and large colony size, with 94% HPDIs ranging from 0.68 to 2.15 (frugivory) and from 1.27 to 1.84 (log_10_ colony size). Membership in Yinpterochiroptera, cave-dwelling, migratory lifestyle, and adult status were also identified as potential risk factors with lower mean posterior densities indicating a weaker association. For paramyxoviruses, the orthorubulaviruses (MuV and PIV2) exhibited a somewhat distinct pattern from MeV and PIV1. Seropositivity against orthorubulaviruses was associated with frugivory, large colony size, and cave-dwelling, similar to flaviviruses, with 94% HPDIs ranging from 1.05 to 2.61 (frugivory), 0.65 to 1.44 (log_10_ colony size), and 0.56-1.49 (cave-dwelling). By contrast, these traits were not apparent risk factors for MeV or PIV1 seropositivity. Similar to the flaviviruses, migratory lifestyle and adult status were identified as weakly associated risk factors for all four paramyxoviruses examined. In contrast to the orthoflavi- and orthorubulaviruses, seropositivity against togaviruses (CHIKV and VEEV) was more prevalent in solitary bats, with a negative relationship to colony size observed (94% HPDIs ranged from -2.47 to -0.19). Models for coronavirus (SARS-CoV, HCoV-229E), pneumovirus (RSV), and phenuivirus (RVFV) indicated that the selected parameters were not strongly predictive of seropositivity, as mean posterior probabilities of all parameters were between -1 and 1, with 94% HPDIs overlapping or near 0. Thus, for coronavirus, pneumovirus, and phenuivirus antigens, it is likely that an insufficient number of seropositive samples were obtained to reliably identify potential associations.

Overall, model B resulted in larger posterior density intervals (greater uncertainty), as effects due to collinear input variables, such as bat family and insectivorous/frugivorous, could not be reliably distinguished. This was particularly noticeable for the orthoflavi- and orthorubulaviruses, such that both membership in Pteropodidae and frugivory were identified as having moderate effect sizes (mean posterior probabilities ranged from 1.02 to 1.29 with respect to membership in Pteropodidae and 0.40 to 0.63 with respect to frugivory), but unlike model A had 94% highest density intervals that overlapped with 0, indicating uncertainty as to the relationship of either trait towards seropositivity. Notably, large colony size remained a clear risk factor for orthoflavi- and orthorubulavirus seropositivity in model B. In contrast to the orthorubulaviruses, MeV did not exhibit an association with Pteropodidae or frugivory, and rather exhibited an association with the Miniopteridae. Regarding togaviruses, both models A and B showed tree-dwelling, small colony size, adult status, and male sex as parameters moderately associated with seropositivity. For SARS-CoV, hCov-229E, and RVFV, the large posterior density intervals in model B precluded assignment of any of the selected parameters as risk factors when accounting for bat family and region_year.

### Potential cross-reactivity of tropical bat sera evaluated using Jaccard similarity

Finally, to investigate potential cross-reactivity in tropical bat sera, we calculated the Jaccard similarity between each pair of viruses that were analyzed (**Fig. 3**). This is important as a serum sample which contains antibodies against an unknown zoonotic virus could theoretically exhibit cross-reactivity against more than one known serogroup, especially given that whole-virus antigen was used. Consistent with our IIFT-based findings using human reference serum (**S2 Table**), three groups suggestive of cross-reactivity were found: the flaviviruses (DENV2, YFV, and WNV), the paramyxoviruses (MuV, PIV2, and, to a lesser degree, PIV1), and the togaviruses (CHIKV and VEEV). High Jaccard similarities between the flavi- and paramyxoviruses were also observed (especially between orthoflavi- and orthorubulaviruses), presumably because of high rates of infection among similar populations (e.g., Pteropodidae bats).

**Fig. 3.**
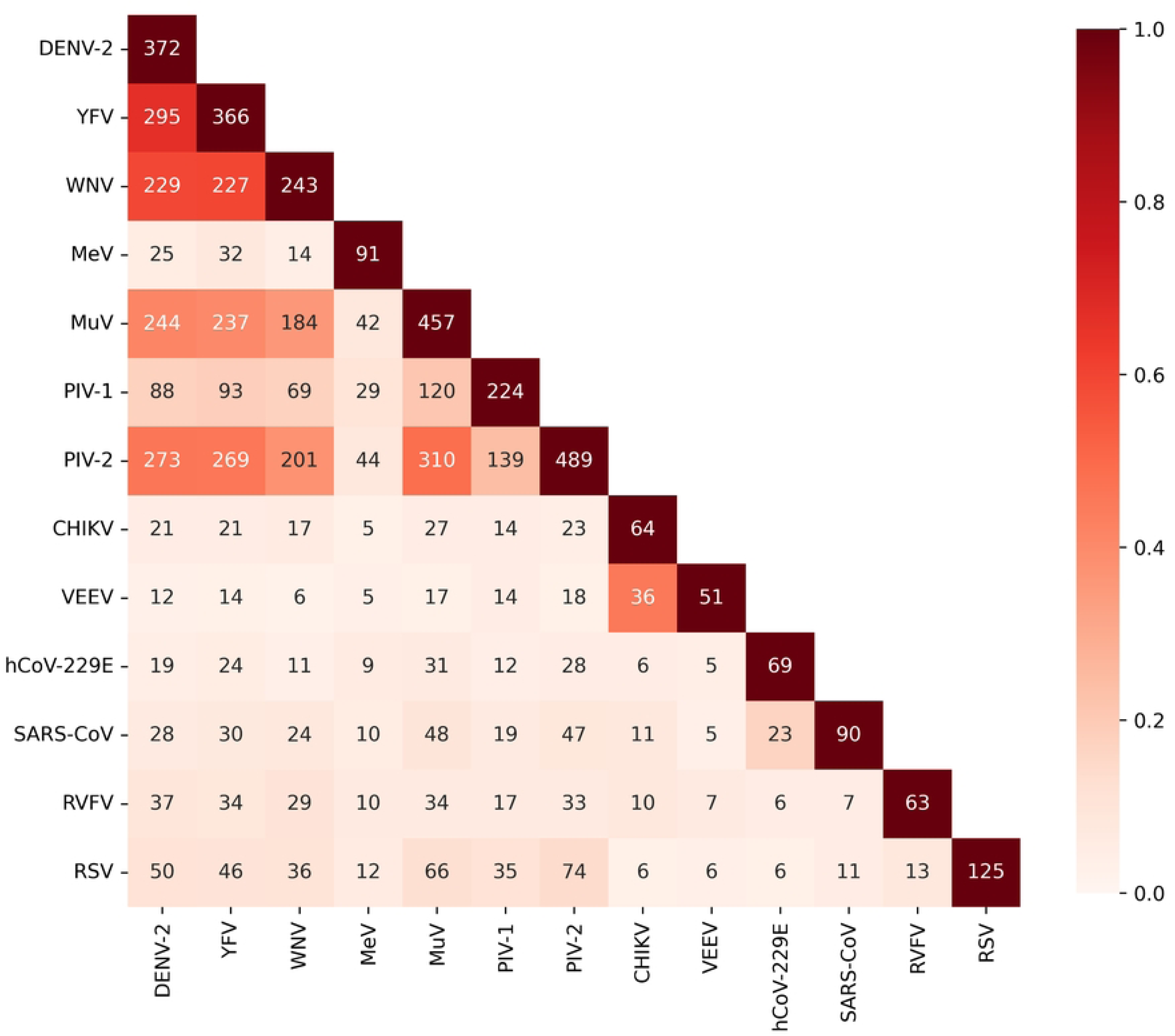
Heatmap showing correlations of seropositivity between tested viruses in tropical bats. Colors represent the Jaccard similarity between seropositivity measurements of pairs of viruses. The coloring scale is shown on the right-hand side. Numbers indicate counts of samples seropositive for both corresponding viruses (indicated on the axes). The Jaccard similarity between measurements of two viruses, *i* and *j*, is defined as follows: 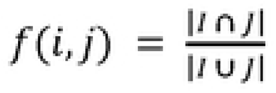, where *I* is the set of samples which were seropositive against virus *i* and *J* is the set of samples which were seropositive against virus *j*. Virus abbreviations were as follows: Dengue virus serotype 2 (DENV-2), yellow fever virus (YFV), West Nile virus (WNV), measles morbillivirus (MeV), mumps virus (MuV), human parainfluenzavirus 1 (PIV-1), human parainfluenzavirus 2 (PIV-2), Chikungunya virus (CHIKV), Venezuelan equine encephalitis virus (VEEV), human coronavirus 229E (HCoV-229E), severe acute respiratory syndrome coronavirus (SARS-CoV), Chikungunya virus (CHIKV), and respiratory syncytial virus (RSV).

## Discussion

The seroprevalences reported in this study provide evidence for the enzootic presence of potentially pathogenic orthoflavi- and orthorubulaviruses in *Eidolon helvum* and *Rousettus aegyptiacus*, as well as evidence for the widespread en-or epizootic presence of all seven virus families examined in both bats and rodents (**Fig. 2; S3 Table**). Notably, other serological surveys of both arbo-[30] and rubulaviruses [31] in African bats are broadly consistent with our study, although the seroprevalence rates we report are somewhat higher, likely due to the use of whole-virus antigen in the mIIFT assay. Determinants of viral seroprevalence are complex and depend on host species, pathogen, and ecology-specific features. For example, despite the relatively high seroprevalence of flaviviruses in African bats, no seropositive samples against flaviviruses were found in European Vespertilionidae (**Fig. 2a**), emphasizing that RNA virus distribution and concomitant risk to humans is highly dependent on local ecological factors. Public health measures designed to minimize the risk of RNA virus spillover should therefore be based on regional risk assessments. The especially high rates of seroprevalence of multiple virus families in *E. helvum* and *R. aegyptiacus* (**Fig. 2a**) may support the implementation of public health measures to reduce human contact with these species. Measures will necessarily be tailored to specific regional requirements, but one successful past example includes interventions in Bangladesh that were shown to be effective at reducing human exposure to potentially Nipah virus-infected *Pteropus* bats [32].

Nonhuman primates are known reservoir and amplification hosts for different flaviviruses and have been shown to play an essential role within the enzootic life cycle of YFV and DENV [15, 33, 34]. Some arthropod-borne flaviviruses cause systemic infection in humans and are capable of replicating in intestinal epithelium and reproductive organs [35–38], raising the possibility that related viruses might circulate among animals via alternative routes of infection. The presence of antibodies against arthropod-borne flaviviruses in bats (**Fig. 2a,c**) [30] suggests a possible role for bats as sylvatic hosts. Additionally, many cross-reactive flaviviruses have been isolated from bats with no known arthropod vector, including Bukalasa, Dakar, Entebbe, Montana *Myotis* leukoencephalitis, and Phnom-Penh bat viruses, raising the possibility of vector-independent circulation among bats [39–42]. Upon their initial discovery, Bukalasa, Dakar, and Entebbe bat viruses were exclusively isolated from the salivary glands of bats [16], suggesting that enzootic transmission of bat flaviviruses via oral route may be a possibility. Many flaviviruses have since demonstrated a capacity to infect other mammals via the fecal or oral routes, or to replicate in the gastrointestinal tract [43–48]. The possibility of frugivory as a risk factor for flavivirus seropositivity among tropical bats (**S6a Fig.**) is consistent with this hypothesis and future studies should further explore the possibility of fecal-oral transmission of flaviviruses, by detecting or isolating virus from contaminated food or feces, which was beyond the scope of this study.

Interestingly, African bats have been previously found to host bat mumps orthorubulavirus, which is cross-reactive with human MuV [49]. Nipah virus (NiV), another bat-borne human pathogen of the paramyxoviruses, exhibits properties consistent with food-borne transmission, increasing the risk for interspecies spillover [50, 51]. Based on the appearance of frugivory as a risk factor for orthorubulavirus seropositivity (**S6b Fig.**), we hypothesize that fruit-borne bat excreta are also a possible route of sylvatic and interspecies rubulavirus transmission and warrant further investigation. For example, a metagenomic, high-throughput sequencing approach to characterize the fecal virome of key bat species, such as *E. helvum* and *R. aegyptiacus* would be practicable with currently available methods and may be valuable for future virus discovery efforts [52]. As revealed by logistic regression model B, however, we cannot presently exclude that taxon-specific factors besides frugivory are responsible for this apparent association with seropositivity (**S7b Fig.**). In contrast to seropositivity against the orthorubulaviruses, which was associated with Pteropodidae bats, seropositivity against MeV was associated with Miniopteridae bats (**S7b Fig.**). Previously, morbilliviruses have been detected in Neotropical *Myotis*, *Desmodus*, and *Phyllostomus* bats [53–55], making it highly plausible that other Yangochiroptera are also susceptible to infection. Our data suggest that Old World Miniopteridae may host an as yet unknown bat morbillivirus.

Consistent with previous reports, this study also provides preliminary evidence for enzootic, communal circulation of orthorubula- and orthoflaviviruses among African megabats, especially *R. aegyptiacus* and *E. helvum* [30, 31, 56–59]. Among tropical bats, logistic regression analysis identified colony-dwelling as a potential risk factor that was predictive of both orthorubula- and orthoflavivirus seropositivity, regardless of the regression model that was employed (**S6 Fig., S7 Fig.**). These data lend support to a previous finding that the exceptionally large colony size of *E. helvum* may contribute to virus spread by exceeding the critical community size, such that a sufficient number of immunologically naïve individuals may continuously be present to support uninterrupted cycles of infection [60].

We also note that togavirus seropositivity (corresponding to mosquito-borne VEEV and CHIKV) was not restricted to insectivorous bats (**Fig. 2**), suggesting that bat predation of insects is not required for sylvatic maintenance of the togaviruses. Additionally, the finding that colony-dwelling bats were less likely to be togavirus seropositive than solitary species (**S6c Fig.**), coupled with a generally lower seroprevalence, might be suggestive of a more sporadic, vector-based mode of transmission, distinct from putative community-based transmission of the orthoflavi- and orthorubulaviruses. Simultaneous PCR sampling of both virus-infected vector and host would be needed to confirm this hypothesis.

Unlike methods involving virus isolation or RNA extraction from field samples, IgG antibodies provide a record of prior infection events and are not limited to a particular body compartment, which contributed to the high sensitivity of our screening approach (**Fig. 2a,b**). While serology-based methods do not definitively identify particular viruses, their specificity is generally defined by viral species delimitation. A degree of cross-reactivity (**S2 Table, Fig. 3**) was a deliberate feature of our study, where our aim was to characterize the general prevalence of infection within larger viral taxa, such as genera. Cross-reactivity was facilitated by the use of whole-virus antigens that necessarily include epitopes conserved across virus genera, such as nonstructural proteins within viral replication complexes. In contrast, in cases where more specific virus detection is required, the specificity of serological detection could be considerably increased by the application of serum-based virus neutralization tests [61].

One significant limitation of this study is its retrospective sampling strategy, which did not allow us to perform large-scale neutralization tests for the different RNA viruses, as only small residual volumes of serum were available. Moreover, as samples were included in this study based on their availability, rather than a pre-meditated sampling strategy, confounding factors make this dataset unsuitable to broadly generalize about the relative seropositivity between bats and rodents or tropical and temperate animals. For example, no African rodents were included as a part of this study, and no temperate regions were included outside of Europe. Future serosurveys should employ sampling strategies which ensure that target species better reflect geographic diversity within each taxonomic order.

In summary, we demonstrate that serological assessment of wildlife is a feasible method for detecting animal RNA viruses with spillover potential. The primary advantage of antibody-based detection methods is the possibility of gathering information about virus prevalence throughout an animal’s lifespan, rather than merely at the point of infection. This approach is especially valuable in studies where detection rates are generally low, or sample sizes are limited. It is important to bear in mind that rates of seropositivity across hosts may not directly correspond to viral prevalence. For example, the longevity of bats (up to 35 years) might contribute to high rates of seropositivity. Another limitation of this study is that the biological factors affecting successful seroconversion in bats remain poorly defined, and differences in immunity among Yango- and Yinpterochiroptera cannot be excluded. Furthermore, antigenic-relatedness is not a perfect correlate of viral pathogenicity, and numerous, difficult-to-quantify factors determine the spillover potential of enzootic viruses [62]. As a technical matter, it was unclear to what degree variation in sample quality (e.g., the inclusion of both transudate as well as serum samples) affected seropositivity rates. Finally, it is important to note that the logistic regression models employed indicate statistical associations of tropical bat characteristics with seropositivity, but do not provide direct evidence that these characteristics are causally responsible for infection. Statistical approaches that are designed to limit multicollinearity, such as model A (**S6 Fig.**), may be more likely to discriminate between various risk factors, but do not necessarily distinguish between underlying variables, such as taxonomic group, which may be critical to infection and therefore affect parameter estimates of the included variables. Model B (**S7 Fig.**), which included additional, collinear variables, increased uncertainty in parameter estimates, but we believe it to represent more accurate estimates for the given variables. Nevertheless, detection of antibodies against pathogenic serotypes can facilitate surveillance of both en- and epizootic viruses of global concern and may provide valuable first indications about how animal pathogens are transmitted among reservoir hosts.

## Acknowledgments

We thank all field workers for assisting in the original sampling, including Jens Jacob for sampling coordination, and Matthias Wenk, Jörg Thiel, Henrike Gregersen, Ulrike M. Rosenfeld and Sabrina Schmidt for providing samples. In addition, we thank Stephan Kallies, Annette Klein, Monika Eschbach-Bludau (all originally University of Bonn Medical Center), Tobias Bleicker, Sebastian Brünink (both Charité – Universitätsmedizin Berlin), Dörte Kaufmann and Mathias Schlegel (both FLI) for excellent technical assistance and support. Samples and essential reagents were contributed by earlier projects funded by the Deutsche Forschungsgemeinschaft, including grants DR772/12-1, DR722/10-1 to CD. The funders had no role in study design, data collection and analysis, decision to publish, or preparation of the manuscript.

## Supporting information Captions

**S1 Table. Colony size and lifespan estimations of tropical bat species.**

**S2 Table. Cross-reactivity testing with N=421 human control sera**

**S3 Table. Complete dataset of tested rodent and bat sera**

**S1 Fig. Directed acyclic graph (DAG) used to inform logistic regression model specification.** Depicted are the proposed parameters used for logistic regression model specification. Arrow direction indicates a known or potential causal relationship between two parameters.

**S2 Fig. Reactivity of goat anti-bat and goat anti-mouse immunoglobulin (Ig) to antibodies of different bat and rodent species by ELISA.** To evaluate antibody detection efficiency across species, two randomly selected serum samples (diluted 1:200) were selected from 16 bat and 12 rodent species and coated in triplicate on Maxisorb ELISA plates. Secondary detection was performed by using a goat anti-bat Ig (Bethyl, Montgomery, AL, USA) or goat-anti-mouse Ig (Dianova, Hamburg, Germany) followed by a donkey anti-goat horseradish peroxidase labeled Ig (Dianova). Optical density (OD) is reported as absorbance at 450 nm minus background absorbance at 630 nm. The dotted lines represent the mean values.

**S3 Fig. Comparison of mIIFT reactivity between bat serum samples with strong and weak ELISA-reactivity.** mIIFT reactivity of serum sample GB687 from *Rousettus aegyptiacus* (a species with high ELISA reactivity, see S2 Fig.) was directly compared to mIIFT-reactivity of serum sample P1018 from *Artibeus jamaicencis* (a species with low ELISA reactivity, see S2 Fig.). Sera were diluted 1:40 and secondary detection was performed with goat anti-bat Ig (1:1000) followed by DyLight488–labeled donkey anti-goat Ig (1:200). P1018 exhibited higher background fluorescence in the DyLight488 channel, but showed specific reactivity against MeV-, MuV-, and PIV2-infected cells as indicated by brightly fluorescent puncta. mIIFT reactivity of bat serum samples was rated in categories: very high reactivity (+++), high reactivity (++), reactivity (+), uncertain reactivity (+/-), and negative (-). Cell nuclei were marked by DAPI staining (blue). All photographs were taken with identical microscope settings and the scale bar represents 25 μm.

**S4 Fig. Mosaic-chip-based indirect immunofluorescence test results of a representative serum sample compared to positive human reference sera.** *R. aegyptiacus* sample GB687 was arbitrarily selected as a representative serum sample exhibiting reactivity against multiple virus antigens. Examples are provided for each of the seven virus families examined in this study: severe acute respiratory syndrome coronavirus (SARS-CoV, *Coronaviridae*); dengue virus serotype 2 (DENV2, *Flaviviridae*); Dobrava-Belgrade orthohantavirus (DOBV, *Hantaviridae*); mumps virus (MuV, *Paramyxoviridae*); Rift Valley fever virus (RVFV, *Phenuiviridae*); respiratory syncytial virus (RSV, *Pneumoviridae*); Chikungunya virus (CHIKV, *Togaviridae*). IIFT results for each serum sample and virus combination were categorized into one of the following: very high reactivity (+++), high reactivity (++), reactivity (+), uncertain reactivity (+/-), and negative (-). Cell nuclei were marked by DAPI staining (blue). All photographs were taken with identical microscope settings and the scale bar represents 25 μm.

**S5 Fig. Evaluation and rating of immunofluorescence signal intensities.** Reactivity of bat serum samples was rated in categories: very high reactivity (+++), high reactivity (++), reactivity (+), uncertain reactivity (+/-), and negative (-). Shown are examples of the mosaic chip-based indirect immunofluorescence test with DENV2-infected cells representing variable fluorescence intensities. All four sera were from *Rousettus aegyptiacus* bats sampled in Gabon. Bat serum specimens were diluted 1:40 in sample buffer, and secondary detection was performed with goat-anti-bat Ig (Bethyl) followed by DyLight488–labeled donkey-anti-goat Ig (Dianova). Cell nuclei were marked by DAPI staining (blue). All photographs were taken with identical microscope settings and the scale bar represents 25 μm.

**S6 Fig. Parameter sizes estimated by multivariable logistic regression using model A.** Posterior distribution of estimated sizes for regression parameters, given as expected change in log odds of detecting seropositivity for seven viruses found in tropical bats. Horizontal black bars beneath distributions indicate the 94% (thin bar) and 50% (thicker and shorter overlapping bar) highest posterior density intervals. Estimated associations of the following sample characteristics with seropositivity are shown: membership in the taxonomic suborder Yangochiroptera (as opposed to Yinpterochiroptera), migratory lifestyle (as opposed to sedentary lifestyle), insectivorous diet (as opposed to frugivorous diet), cave-dwelling (as opposed to tree-dwelling), Log_10_ colony size (shown for 5 units on the log_10_ scale), adult (as opposed to juvenile), and male (as opposed to female). Results shown for **(a)** flaviviruses, **(b)** paramyxoviruses, **(c)** togaviruses, **(d)** coronaviruses, **(e)** a pneumovirus, and **(f)** a phenuivirus.

**S7 Fig. Parameter sizes estimated by multivariable logistic regression using model B.** Posterior distribution of estimated sizes for regression parameters, given as expected change in log odds of detecting seropositivity for seven viruses found in tropical bats. Horizontal black bars beneath distributions indicate the 94% (thin bar) and 50% (thicker and shorter overlapping bar) highest posterior density intervals. Estimated associations of the following sample characteristics with seropositivity are shown: membership in the taxonomic suborder Yangochiroptera (as opposed to Yinpterochiroptera), migratory lifestyle (as opposed to sedentary lifestyle), insectivorous diet (as opposed to frugivorous diet), cave-dwelling (as opposed to tree-dwelling), Log_10_ colony size (shown for 5 units on the log10 scale), adult (as opposed to juvenile), male (as opposed to female), bat family, and year_region, a variable comprising both the region and year of sampling. Results shown for **(a)** flaviviruses, **(b)** paramyxoviruses, **(c)** togaviruses, **(d)** coronaviruses, and **(e)** a phenuivirus.

## Author contributions

Conceptualization, J.E., J.P., M.A.M., C.D.; Methodology, J.E., J.P., L.M., T.V., V.M.C., T.B., E.L., B.M.K., S.Z., K.F., F.G.-R., P.V., S.O., C.B.E.M.R., R.G.U., E.M.L, J.F.D, T.J., C.D., M.A.M.; Software, J.E., L.M., T.V., T.J.; Validation, J.E., J.P.; Formal Analysis, J.E., J.P., L.M., V.M.C., M.A.M.; Investigation, J.E., J.P., M.A.M., C.D.; Resources, V.M.C., T.B., E.L., B.M.K., S.Z., K.F., F.G.-R., V.C., G.D.M., J.S.-C., P.V., S.O., M.T., C.B.E.M.R., R.G.U., E.M.L, J.F.D, T.J., C.D., M.A.M.; Data Curation, J.E., J.P., L.M., T.V., V.M.C., T.B., E.L., B.M.K., S.Z., K.F., F.G.-R., P.V., S.O., C.B.E.M.R., R.G.U., E.M.L, J.F.D, T.J. C.D., M.A.M.; Writing – Original Draft Preparation, J.E., J.P., L.M., M.A.M.; Writing – Review & Editing, J.E., J.P., J.F.D., C.D., M.A.M.; Visualization, J.E., J.P., L.M., V.M.C., M.A.M.; Supervision, C.D., M.A.M.; Project Administration, M.A.M.; Funding Acquisition, C.D. All authors reviewed the manuscript.

## Ethics Statement

Trapping of rodents in Germany was conducted in the framework of Hantavirus monitoring activities coordinated by author Rainer Ulrich under the permit numbers LALLF M-V/TSD/7221.3-2.1-030/09, TH 15-107/09, and BW 35-9185.82/0261. The collection of rodents in Thuringia (TH), Baden-Wuerttemberg (BW) and Mecklenburg-Western Pomerania (MWP) was commissioned and funded by the Federal Environment Agency (UBA) within the Environment Research Plan of the German Federal Ministry for the Environment, Nature Conservation, Building and Nuclear Safety (BMUB, UFO-Plan FKZ: 3709 41 401). All procedures involving animals were covered by relevant legislation (permits for BW: Regierungspräsidium Stuttgart 35-9185.82/0261, MWP: Landesamt für Landwirtschaft, Lebensmittelsicherheit und Fischerei Mecklenburg-Vorpommern 7221.3-030/09, THR: Thüringer Landesamt für Lebensmittelsicherheit und Verbraucherschutz 22-2684-04-15-107/09). Additional animals were provided by forestry institutions and pest control in Thuringia, Brandenburg and Baden-Wuerttemberg, which caught and sacrificed the animals during their official duties without the necessity of further permits. Rodent sampling in The Netherlands was licensed by the Dutch Animal Ethics Committee (DEC) under permit numbers 200700119, 200800113, and 200800053 to author Chantal Reusken. Rodent sampling in Gabon was supervised by author Eric Leroy and licensed by the Ministry of Water and Forest, statement 003/MEF/SG/DGEF/DFC from 2011. Rodent sampling in Thailand was granted by the Agricultural Zoology Research Group of the Department of Agriculture in Thailand (permit no. KU./14-182). Sampling and capture of bats in Ghana and Panama, as well as sample transfers, were supervised by author Christian Drosten and done under wildlife permits and ethics clearances: Panama (Research-Permit STRI 2563 (PI)-Veronika Cottontail IACUC 100316-1001-18/Research-Permit ANAM: SE/A-68-11/Ethics-Permit: IACUC 100316-1001-18/Export Permits: SEX/A-30-11, SEX/A-55-11, SEX/A-81-10, SEX-A-26-10); Ghana (Research Permit: 2008–2010 (A04957)/Ethics-Permit: CHRPE49/09/Export-Permit: State Agreement between Ghana and Hamburg (BNI)); Gabon (Ethics-Permit: 00021/ MEFEPA/SG/DGEF/DFC); Germany (Ethics-Permit: LANU 314/5327.74.1.6).

